# Delineating feedback activity in the MAPK and AKT pathways using feedback-enabled Inference of Signaling Activity

**DOI:** 10.1101/268359

**Authors:** Bram Thijssen, Katarzyna Jastrzebski, Roderick L. Beijersbergen, Lodewyk F.A. Wessels

## Abstract

An important aspect of cellular signaling networks is the existence of feedback mechanisms. However, due to the complexity of signaling networks, as well as the presence of multiple interrelated feedback events, it can be difficult to identify which signaling routes are active in any particular context. We have previously shown that Inference of Signaling Activity (ISA) can be a useful method to study steady-state oncogenic signaling across different cell lines and inhibitor treatments. However, ISA did not explicitly include feedback signaling events. Incorporating feedback will increase the complexity and computational cost of the model, and more data is likely to be needed to infer feedback activities. Here, we developed feedback-ISA (f-ISA), an extension of the ISA modeling approach which incorporates feedback signaling events. It also includes integrated batch correction in order to fit the models to multiple, independent datasets simultaneously. We find that the identifiability of feedback activities can be counter-intuitive, which shows the importance of analyzing the full, joint uncertainty in model parameters. By iteratively adapting the model and including multiple datasets, including both steady state and intervention data, we constructed a model that can explain a large part of the phosphorylation levels of several signaling molecules in the MAPK and AKT pathways, across many breast cancer cell lines and across various conditions. The resulting model delineates which routes in the signaling network are likely to be active in each cell line and condition, given all of the data. Additionally, such models can indicate whether datasets agree with each other, and identify which parts of the data cannot be explained, thereby highlighting gaps in the current knowledge. We conclude that this modeling approach can be useful to quantitatively understand how complex cellular signaling networks behave across different cell lines and conditions.

## Introduction

The mitogen-activated protein kinase (MAPK) and AKT signaling pathways are two central regulatory mechanisms which are often deregulated in cancer cells. Many inhibitors which block key kinases in these signaling pathways have been developed as potential cancer therapies. These kinase inhibitors can be very potent anticancer drugs, but cancer cells are often intrinsically resistant or develop resistance over time [1]. As a result, patients have a highly variable response to kinase inhibitors, and this variability is also seen in cell lines [2–4]. To understand this variability in response, and develop effective and selective (combination) therapies, a detailed quantitative understanding of cellular regulatory networks would be highly useful.

Signaling networks are complex, with many interrelated signaling events. One of the important features of cellular signaling networks is the existence of feedback mechanisms. These feedback loops can, for example, provide robustness to a signaling network or modulate the sensitivity to external inputs [5]. To date, many details of numerous signaling networks have been discovered, including various feedback events. In the MAPK pathway, an important feedback mechanism is the inactivation of RAF by ERK [6,7]. In the AKT pathway, two feedback mechanisms involve the down-regulation of the insulin receptor substrate IRS1 [8], and the modulation of SIN1 activity [9,10], a component of the MTORC2 complex. Nevertheless, several aspects of these feedback loops remain unclear.

For instance, in the case of SIN1, there has been debate about the regulation and function of its phosphorylation sites, in particular T86 and T398. Liu *et al* [9] have shown that both S6K and AKT can phosphorylate SIN1, and that this phosphorylation suppresses MTORC2-mediated phosphorylation of AKT. Conversely, Humphrey *et al* [11] have shown that AKT phosphorylates SIN1 and that this stimulates MTORC2 instead. Yang *et al* [10] further investigated this in more cell lines and conditions, arguing that AKT rather than S6K is the major SIN1 kinase and that the feedback is positive, in line with Humphrey *et al*.

For IRS1, one aspect which is unclear is the phosphorylation of S312 (in human IRS1). Note that this site corresponds to mouse S307, while the human S307 constitutes yet another phosphorylation site on IRS1. Signaling by IRS1 is regulated by multiple phosphorylation events [12], where serine/threonine phosphorylation inhibits its activity while tyrosine phosphorylation activates it. For S312, it is known that JNK (among other kinases) phosphorylates this site [13], but it has also been shown that this phosphorylation is MTORC1-dependent [14]. S6K phosphorylates IRS1 on several other sites, including S307 [15] and S270 [16]. Given the MTORC1-dependence of IRS1 S312 phosphorylation, it is sometimes assumed that S312 phosphorylation is also S6K-dependent. Indeed the databases Uniprot [17]and PhosphoSitePlus [18] currently list S6K as kinase for IRS1 S312 phosphorylation (putatively in the case of PhosphoSitePlus), even though to the best of our knowledge there is no direct evidence for this. Rather, it has been reported as unlikely [15], and S312 is also not part of an S6K target motif. MTORC1 may also affect S312 phosphorylation through an effect on protein phosphatase PP2A [19] or indirectly through dependencies between phosphorylation sites. In addition to these details, it is unclear in which contexts feedback through IRS1 -- whether mediated by MTORC1, JNK, S6K or yet other regulators -- is important. For example, feedback to IRS1 is involved in insulin resistance [8], and mediates re-activation of the AKT pathway after rapamycin treatment [20], but seems not to be involved in re-activation of AKT after AKT-inhibitor treatment [21,22].

Computational models can be used to better understand these complex regulatory networks. Different modeling frameworks have been used to quantitatively study oncogenic cellular signaling pathways, including dynamic models [23,24] and steady state models [25–27]. Dynamic models can describe detailed kinetics of a system, but they are costly to simulate, especially when the full parameter uncertainty is analyzed [28–30]. The computational cost is further exacerbated when multiple datasets are included, resulting in many model conditions which have to be evaluated for each parameter value. Logic models allow significantly larger models to be evaluated [31,32], but it is more difficult to model quantitative differences between cell lines and conditions in such a framework.

We have previously developed Inference of Signaling Activity (ISA), a steady state modeling approach to study signaling activities across cell lines along with different inhibitor treatments [27]. One major assumption in ISA is the absence of feedback signaling, which allowed fast model evaluations and hence relatively large signaling models. Models without feedback can give good fits to drug response cell viability data [27], arguing that feedback events are not crucial to describe differences in the relative viability between cell lines after 72 hour drug treatment. However, it is clear that feedback signaling events are a major component of cellular signaling, and are important to consider in many situations. We therefore set out to expand ISA with feedback signaling events, to explore whether the method could also be used to infer signaling activities in models that include feedback events.

## Methods

Below, we outline the substantial changes we have made to ISA [27], resulting in feedback-ISA (f-ISA), an approach capable of modelling feedback mechanisms in signaling pathways across different cell lines and conditions.

### Model structure

In ISA, the activities of signaling molecules are modeled by a continuous latent variable *x*_*i*_ that can take values between 0 and 1. This signaling activity is denoted by *x*_*i*_ where *i* indexes the signaling molecule. This activity is a function of the upstream signaling nodes, as well as a basal activity, the expression of the signaling molecule itself and any kinase inhibitors that may be present. In the original ISA, the activity was restricted between 0 and 1 using a clamping function. In order to accommodate feedback loops, we now change the activation function to a logistic function. Specifically, we calculate the signaling activity as

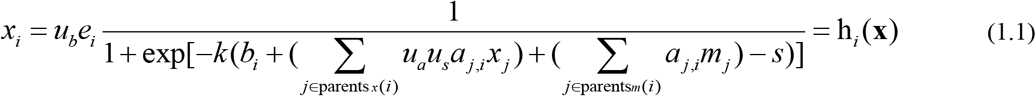

Here *u_a_*, *u*_*b*_ and *u*_*s*_ are the kinase inhibitor effects (defined in more detail later), *e*_*i*_ is the expression of signaling molecule *i*, *b*_*i*_ is the basal activity of the signaling molecule *i*, *a*_*j,i*_ is the strength of signaling from molecule *j* to molecule *i*, *m*_*j*_ is a binary variable denoting whether mutation *j* is present, while *k* and *s* are the constant steepness and inflection point of the logistic function. The logistic function is more expensive to compute than a clamping function, but in contrast to a clamping function, the logistic function is smooth (i.e. continuously differentiable), which simplifies the process of solving the systems of equations with feedbacks. To keep the number of parameters manageable, we fix both the steepness *k* and the inflection point *s* to a set value. The steepness is set to 9.19024 such that the activity is 0.01 and 0.99 when the total input is 0 and 1 respectively, and the inflection point is set to 0.5. An example of an activity calculation is shown in Figure 1B.

**Figure 1:**
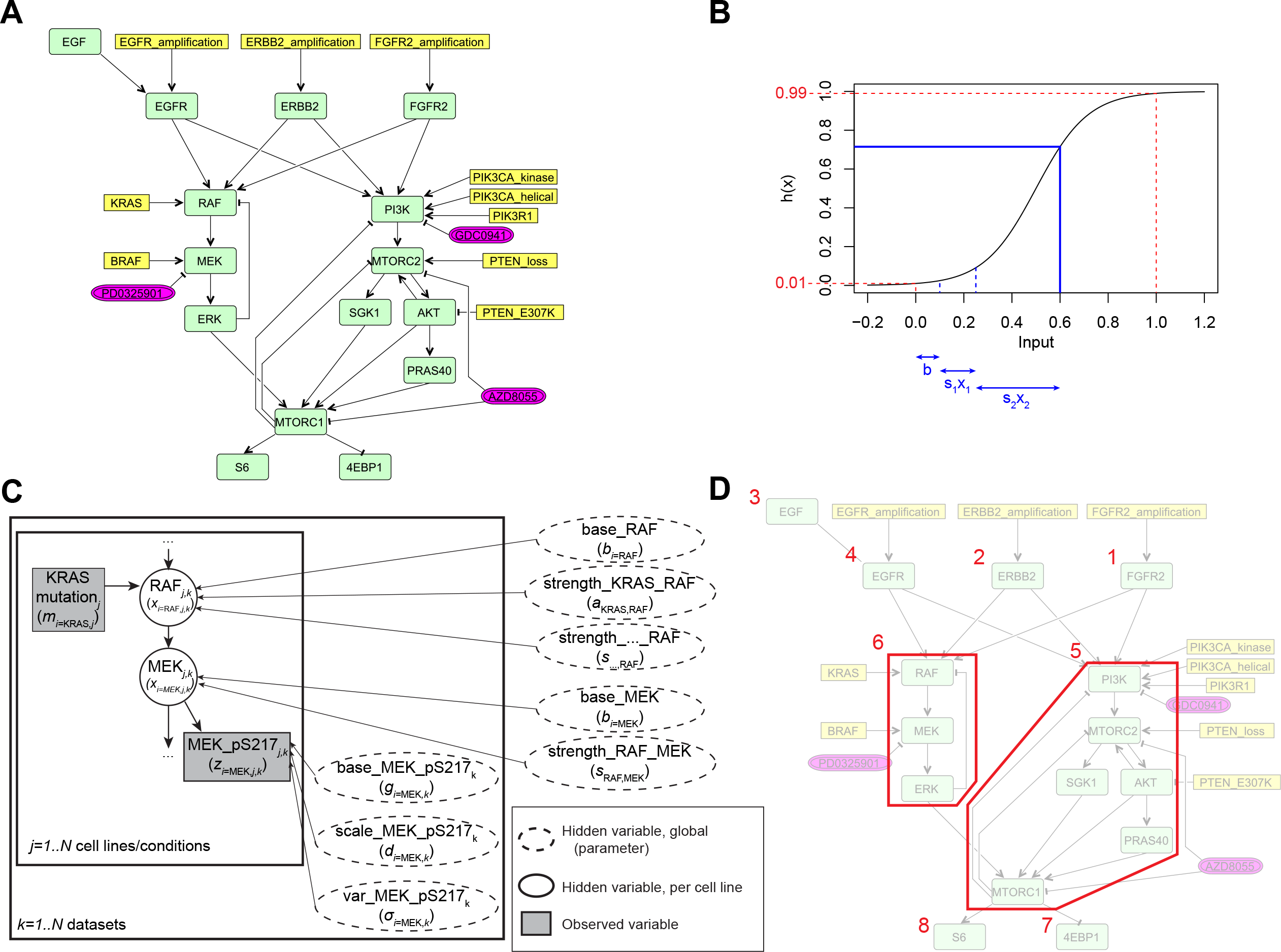
Signaling model and system decomposition. (A) The signaling model of MAPK and AKT signaling in breast cancer constructed for the purpose of these studies. It is based on a reduced, joint-drug model of Jastrzebski *et al* [27]. The model is depicted in Activity Flow format of the Systems Biology Graphical Notation. Yellow boxes indicate genetic events including mutations and copy number aberrations, green nodes are the signaling molecules, and purple nodes are the kinase inhibitors. (B) The activation function used to calculate the activity of a signaling molecule from its input. The red lines indicate the constraints which were taken to determine the fixed steepness and inflection point of the logistic function. The blue lines give an example; in this case the signaling molecule has two upstream inputs (*x*_1_ and *x*_2_), which are multiplied by the respective signal strengths (*s*_1_ and *s*_2_) and summed together with a basal activity (*b*) to give a final signaling activity of approximately 0.7. (C) A small part of the model shown in template notation, illustrating the latent variable structure and the integrated batch correction. The text names indicate the names in the code while the symbols in brackets correspond to the equations in the Methods section. Each dataset (indexed by *k*) has multiple cell lines or conditions (indexed by *j*). The signaling activities are unique for each condition. The likelihood of the data depends on two batch correction variables (base and scale) as well as the variance; these variables are specific for each dataset. The signaling strengths and base activities are global parameters, which are shared by all datasets and all conditions. (D) Decomposition of the model into smaller modules that have to be solved as systems of equations (the red boxes; one system of 3 nodes and one system of 6 nodes), and a topological ordering (the red numbers) giving the order in which to calculate the nodes and systems.

Without feedback events, the signaling activities can be calculated from the upstream molecules downwards. However, by including feedback events, the equations for the signaling activities become coupled, and as a result the activities have to be calculated by solving a system of nonlinear equations. We use the Newton-Raphson method for this; that is, we solve the equation

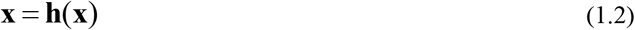

by iterating through

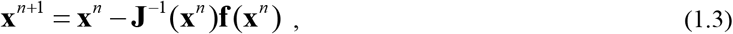

where **x**^*n*^ represents the solution at the *n*-th iteration, and

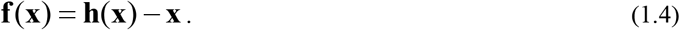

The Jacobian matrix **J** is given by

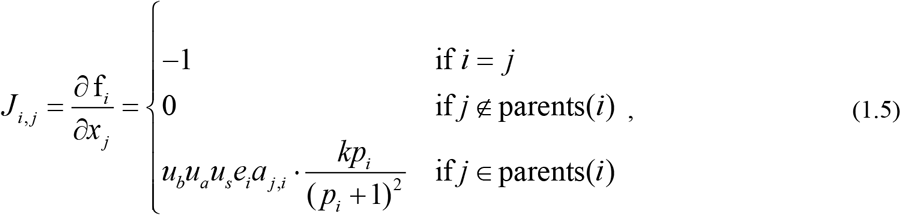

with

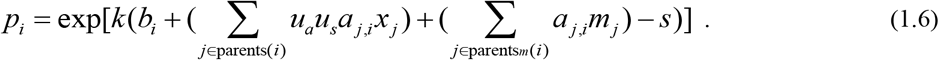

If the system of equations contains four or fewer signaling molecules, Equation 1.3 is most efficiently calculated by taking the inverse of the Jacobian directly. If the system contains more than four signaling molecules, it is more efficient to use LU-decomposition to solve Equation 1.3 instead. We stop the Newton-Raphson iteration when the last change in **x** is less than 10^−5^ in each direction.

Although it is possible to calculate the entire model with Equations 1.2 and 1.3, the LU-decomposition scales cubically in the number of signaling molecules. It would therefore be beneficial to decrease the system size as much as possible. The system of equations is not fully coupled however. Some signaling molecules are not involved in any feedback loop, but are only upstream or downstream of other molecules. The activities of these molecules can be calculated directly, without having to solve a system of equations. Additionally, some signaling molecules which are affected by a feedback loop, may not be affected by another loop. We can therefore decrease the size of the systems to be solved by decomposing the model into smaller systems composed of signaling molecules which are coupled by a feedback signaling loop. To identify which parts need to be solved as a system of equations, we use Tarjan’s strong connectivity algorithm [33] as a preprocessing step. This algorithm identifies all the strongly connected components of a directed graph, as well as providing a topological ordering of the graph. To calculate the signaling activities, we then iterate over the topological ordering, and calculate the activities either directly (for connected components of size 1, i.e. individual signaling molecules that are not part of any feedback system) or by solving the corresponding system of equations (for connected components with size larger than 1). Figure 1C illustrates the decomposition of a model into individual molecules and systems.

Note that the signaling activity vector **x** is a function of the mutations, **m**, protein expression, **e**, and the presence of any inhibitors, *u*, (all of which can change between cell lines and conditions), as well as a number of parameters (which remain constant between cell lines and conditions). As such, the model is constrained to explain all observed signaling activities from the mutation and protein expression patterns. This is a fundamentally different approach from estimating signal strengths separately for each cell line and condition, as done for example by [24]. Rather than inferring cell-line specific signaling strengths, we attempt to infer signal strengths that can fit the data across all cell lines and conditions.

Kinase inhibitors in the model can be of two types: they can either inhibit the activity of their target, or they can inhibit the activation of the target. The type of the inhibitor is derived from the literature. For example, the AKT inhibitor GSK690693 is an ATP-competitive inhibitor which inhibits the activity of AKT, while the AKT inhibitor MK2206 is an allosteric inhibitor which inhibits the activation of AKT. For activity-inhibiting drugs we set *u*_*a*_=*u* in equation 1.2 (for the specific signaling link that is inhibited), and for activation-inhibiting drugs we set *u*_*b*_=*u* (for the specific target that is inhibited). Note that *u*_*a*_ affects the activity of the children of the target, whereas *u*_*b*_ affects the activity of the target itself. The type of the inhibitor is specified in the SBML annotation (described later). Regardless of whether *u*_*a*_ or *u*_*b*_ is used, the inhibitory effect is calculated as

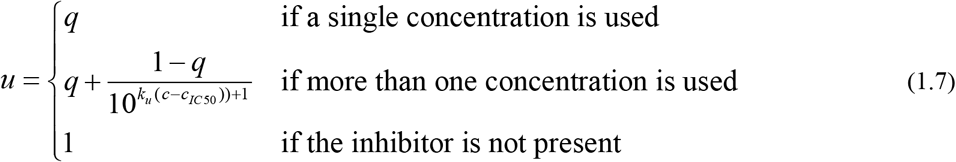

where *q* is the maximal inhibitory effect (or the inhibitory effect at the particular concentration used if there is only one concentration), *k*_*u*_ is the steepness of the inhibitory curve, *c* is the concentration and *c*_*IC50*_ is the 50% inhibitory concentration. In contrast to the activation function in equation 1.1, the steepness and inflection point of the kinase inhibition curves (i.e., *k*_*u*_ and *c*_*IC50*_) are included as free parameters to be inferred. Finally an inhibitor may increase the susceptibility of its target to *incoming* signals, a process which has been named “inhibitor hijacking” [21]. The variable *u*_*s*_ is included in Equation 1.1 to reflect this effect. This parameter is only used if the drug is specified to alter the susceptibility of its target, and we use it specifically to allow ATP-competitive AKT inhibitors to alter the susceptibility of AKT phosphorylation by MTORC2.

### Likelihood function

Jastrzebski *et al* [27] used protein phosphorylation data as well as untreated proliferation rates and relative viability upon kinase inhibitor treatment to infer the signaling activities. However, since f-ISA is computationally more expensive than the original ISA framework, we only use protein phosphorylation data for the inference here. Including cell viability data, in particular dose response curves, would result in too many model conditions as well as additional parameters to be estimated. Mutation data, copy number data and total protein expression data are still used as before [27], by directly setting the corresponding variable (*m*_*i*_ or *e*_*i*_) to the observed value. The structure of a small part of a model is shown in template notation in Figure 1D.

The likelihood of an observed protein phosphorylation measurement is defined as

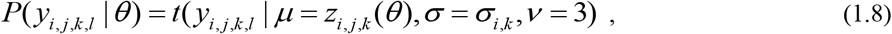

where *Θ* is the vector containing all model parameters, *y*_*i,j,k,l*_ is the measurement data for observed variable *i*, cell line *j*, dataset *k* and replicate *l*, *z*_*i,j,k*_ is the modeled variable (defined further below in Equation 1.9) and *σ*_*i,k*_ is the variance of observed variable *y*_*i*_ in dataset *k*. The variance *σ*_*i,k*_ for observed variable *i* is shared by all cell lines and biological replicates, but is specific for each dataset *k*.

The modeled variable *z* is defined as

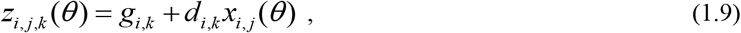

where *g*_*i,k*_ is the background signal generated by aspecific binding of the antibody and *d*_*i,k*_ is a scaling factor to account for differences between datasets (recall that *k* indexes the dataset). If a particular phosphorylation is only measured by one dataset, then *d*_*i,k*_ is set to 1, and if the phosphorylation is measured by more than one dataset, then *d*_*i,k*_ is included as a free parameter to be inferred, for every dataset. *g*_*i,k*_ is always included as free parameter to be inferred for every epitope and dataset, regardless of whether the epitope is uniquely measured by only one dataset. Finally, we assume independence between the epitopes, datasets, cell lines and replicates, and the total likelihood is calculated as a product of the likelihoods of the individual data points.

Although the likelihood function includes a scaling variable to perform batch correction, it is useful to have the measurements on approximately the same scale between datasets before running the inference. Therefore, as a preprocessing step, the measured phosphorylation levels are reduced to [0,1] by dividing the measurements of each epitope by the maximum value observed for that epitope in any cell line or condition in that dataset, if that value is larger than 1. With the RPPA quantification procedure that was used (SuperCurve [34]), maximum observed values below 1 indicate that for that epitope, all conditions in that dataset have low phosphorylation levels, relative to the set of standard lysates. We therefore do not *increase* the signal to 1, to prevent amplifying experimental noise. With all measurement values in the range [0,1], the prior for the scaling parameters *d*_*i,k*_ can then be set to a small uniform distribution, between 0 and 2, which reduces the parameter space that has to be searched.

### Parameter priors

The model contains several different types of parameters that are inferred from the data. Table 1 lists these parameters, along with the prior distribution that is used for each of them.

**Table 1.**
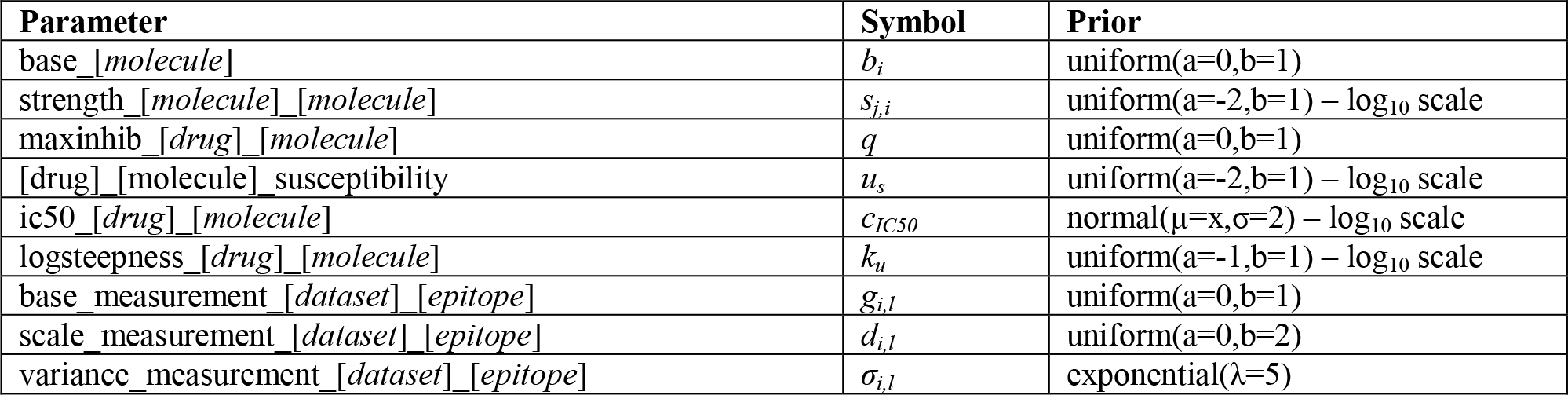
Model parameters that are inferred from the data.

| Parameter | Symbol | Prior |
| --- | --- | --- |
| base_<sub>[molecule]</sub> | $b_i$ | uniform(a=0,b=1) |
| strength_<sub>[molecule]</sub>_<sub>[molecule]</sub> | $s_{j,i}$ | uniform(a=-2,b=1) – log<sub>10</sub> scale |
| maxinhib_<sub>[drug]</sub>_<sub>[molecule]</sub> | $q$ | uniform(a=0,b=1) |
| [drug]_<sub>[molecule]</sub> susceptibility | $u_s$ | uniform(a=-2,b=1) – log<sub>10</sub> scale |
| ic50_<sub>[drug]</sub>_<sub>[molecule]</sub> | $c_{IC50}$ | normal($\mu=x, \sigma=2$) – log<sub>10</sub> scale |
| logsteepness_<sub>[drug]</sub>_<sub>[molecule]</sub> | $k_u$ | uniform(a=-1,b=1) – log<sub>10</sub> scale |
| base_measurement_<sub>[dataset]</sub>_<sub>[epitope]</sub> | $g_{i,l}$ | uniform(a=0,b=1) |
| scale_measurement_<sub>[dataset]</sub>_<sub>[epitope]</sub> | $d_{i,l}$ | uniform(a=0,b=2) |
| variance_measurement_<sub>[dataset]</sub>_<sub>[epitope]</sub> | $\sigma_{i,l}$ | exponential($\lambda=5$) |

The strength parameter for reactions with a known influence sign is inferred on a logarithmic scale. This is done to give equal prior weight to weak interactions (strength 0.01-0.1), interactions with intermediate strength (0.1-1) and strong amplifying interactions (1-10). If this range from 0-10 was not log-transformed and a uniform distribution would still be used, most of the prior weight would be on amplifying signals, and all signaling activities would be heavily saturated *a priori*, especially in a signaling cascade. When the sign of the influence is not known, that is, when the signal could be either activating or inhibiting, then the prior is set on a regular scale from −2 to 2.

For drug inhibition, the drug can be specified to either use a single drug concentration, giving only a single parameter for the inhibition at that concentration, or to use multiple drug concentrations, giving three parameters: the maximum inhibition, the 50% inhibitory concentration (IC_50_) and the steepness. The prior for the 50% inhibitory concentration is centered on the concentration at which it inhibits its target for 50%, as determined by *in vitro* inhibition experiments described in the literature.

### Model structure specification

The signaling graph of the model is constructed in CellDesigner [35], version 4.4. Using this tool, the model is specified in Systems Biology Graphical Notation as an Activity Flow diagram [36]. That is, rather than specifying precise molecule states and reactions, only the abstract influences between signaling molecules are specified, which fits with the modeling paradigm of ISA. The model is stored as an SBML file with CellDesigner extension annotation. This extension annotation allows specification of the types of the signaling molecules (i.e. whether a node is a signaling molecule, mutation or drug) as well as specification of the signaling influences and the types of drug inhibition.

### Inference

To infer the posterior distribution of the parameters, we use the BCM software package [37]. To incorporate f-ISA in BCM, we developed the tool `sbmlpdinf`, which can read the activity flow diagram described in the SBML/CellDesigner file, as well as two XML files specifying the likelihood and the prior. All data is stored in a single NetCDF4 file. The likelihood file specifies which measurement in the data file corresponds to which node in the model, as well as all the model conditions, such as the presence of a particular kinase inhibitor. The model simulation code was written in C++ and made thread-safe for use in the parallelized inference algorithms of BCM. The largest model presented here, with 18 signaling molecules and two feedback systems, could be evaluated at approximately 1.1 million model evaluations per second using 18 threads on an Intel Xeon E5-2697 v4 processor.

We sampled the posterior distribution using feedback-optimized, parallel tempered MCMC [38] with automated parameter blocking [39]. The marginal likelihood was calculated using thermodynamic integration [40]. The temperature schedule was optimized twice, which also served as burn-in period. After this optimization, 1,000 samples were generated from the posterior (after subsampling), with the amount of subsampling chosen such that the autocorrelation was negligible and at least 100 roundtrips from prior to posterior were performed. Specifically, when one dataset is included in the inference we use 24 parallel chains and run the inference with a subsampling of 1 in 2,000 and probability of choosing a temperature swap move of 0.9. Each MCMC move constitutes updating every parameter or parameter block once. When more than one dataset is included we increased the number of chains to 36 and use a subsampling of 1 in 4,000. Sample traces are provided in Supplementary File 1. Despite the high performance implementation, inference with a single dataset and a medium-sized model takes several hours, and the largest inference presented here, which included 108 model conditions and 139 free parameters, required approximately 48 hours to run with the aforementioned processor.

### Quantifying reduction in uncertainty

To quantify the reduction in uncertainty, we used the Occam factor introduced by MacKay [41], which quantifies the change in volume of the parameter space that is accessible from the posterior compared to the prior. In other words, it measures which fraction of the prior parameter space is consistent with the data. MacKay used a Gaussian approximation to calculate the Occam factor; but since we evaluated the marginal likelihood as part of the inference here, we can calculate the Occam factor directly by

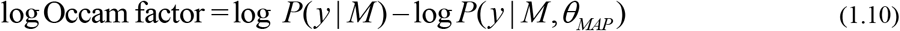

where *P*(*y*|*M*) is the marginal likelihood of the data given the model and *P*(*y*|*M,*θ*_MAP_*) is the likelihood of the data at the maximum a posteriori value of the parameters.

### Cell lines

All cell lines used have been described, including their culture conditions, in Supplementary Table 1 of Jastrzebski *et al* [27].

### Measurement of on-treatment phosphorylation

Cell lines were seeded in 60 mm dishes (BT549 at 4×10^5^ cells/dish; HCC1954 at 8×10^5^ cells/dish; MCF7 at 4×10^5^ cells/dish; MM231 at 8×10^5^ cells/dish; MM453 at 1×10^6^ cells/dish; MM468 at 1×10^6^ cells/dish; SKBR3 at 6×10^5^ cells/dish; T47D at 8×10^5^ cells/dish). Following 24 h of incubation, all but the exponentially growing cells were serum starved in unsupplemented base medium containing penicillin/streptomycin (Gibco) for a further 24 h. Cells were then treated with either DMSO vehicle control or one of the following three inhibitors - PD0325901 at 50 nM, GDC0941 at 10 μM or AZD8055 at 1 μM – for 30 min, after which stimulation with 10 ng/ml EGF, where indicated, was carried out for a further 20 min. Cells were then placed on ice, washed with ice-cold PBS, and lysed in 150 μl/dish of RIPA buffer (20 mM Tris-HCl, pH 8, 150 mM NaCl, 1% NP40, 0.5% sodium deoxycholate, 0.1% SDS) supplemented with cOmplete protease and phosSTOP phosphatase inhibitor cocktails (Roche). Lysates were cleared by centrifugation at 4°C and 20,800 × g, and protein concentration determined using the Pierce BCA protein assay (Thermo Fisher Scientific). The supernatant was normalized to 1 μg/μl with RIPA buffer and supplemented with SDS sample buffer to a final concentration of 62.5 mM Tris-HCl pH 6.8, 10% glycerol, 2% SDS, 2.5% (v/v) 2-mercaptoethanol. Samples were further assayed at the MD Anderson Cancer Center RPPA Core Facility, as outlined previously [27].

### IRS1 disruption experiment

Cell lines were seeded in 6-well plates (BT549 at 4×10^5^ cells/well; HCC1954 at 2×10^5^ cells/well), and following a 48 h incubation, treated with either DMSO vehicle control or 5 μM NT157 for a further 24 h. They were then treated for a further 1 h with either DMSO vehicle control, 1 μM GSK-690693 or 5 μM PF-4708671. Cells were then lysed and protein concentration determined as outlined above. Twenty μg of protein were supplemented with Novex^®^ LDS Sample Buffer and Sample Reducing Agent, heated at 70°C for 10 min and separated on 4-12% gradient gels (Thermo Fisher Scientific). Separated proteins were transferred onto Immobilon-P PVDF membranes (Merck Millipore) using a Trans-Blot^®^ system (Bio-Rad). Blocking was performed in TBS supplemented with 0.1% Tween and 3% BSA (TBS-TB) for 1 h at room temperature, followed by overnight immunoblotting at 4°C with the following primary antibodies: IRS1 (MERCK/Millipore 06-248); pIRS1-S312 (Cell Signaling Technology 2381); AKT (Cell Signaling Technology 9272); pAKT-S473 (Cell Signaling Technology 4060); SIN1 (Cell Signaling Technology 12860); pSIN1-T86 (Cell Signaling Technology 14716); PRAS40 (Cell Signaling Technology 2610); pPRAS40-T246 (Cell Signaling Technology 2997); S6 (Cell Signaling Technology 2217); pS6 (S235/236) (Cell Signaling Technology 2211); Vinculin (Sigma V9131). Membranes were then washed with TBS supplemented with 0.1% Tween (TBS-T) and probed with secondary goat anti-mouse or anti-rabbit HRP-conjugated antibodies (Bio-Rad) diluted in TBS-TB for 2 h at room temperature. Finally, membranes were washed in TBS-T, an ECL reaction was carried out using the Clarity^TM^ Western ECL Substrate (Bio-Rad) and the signal detected using a ChemiDoc Touch instrument (Bio-Rad). Quantification of the exposures was performed using ImageQuant software (Bio-Rad).

### External data

In addition to the data we generated here, we included a part of the dataset provided by Korkola *et al* [42] in the inference. They performed RPPA measurements of 15 breast cancer cell lines, treated with either lapatinib (250 nM), GSK690693 (250 nM), a combination of the two, or a DMSO control, at 8 time points from 30 minutes to 72 hours. Since we focus here on fast-acting post-translational feedback here, we selected the 1-hour time point from their data, which most closely matched the 50-minute treatment of our on-treatment phosphorylation measurements. We used only the 9 cell lines which overlap with our cell line panel, since we have mutation and copy number data available for these cell lines. To have the measurements on the same scale, we reverse-log-transformed the data and divided by the maximum value for each epitope as described above.

## Results

### Constructing a model of steady state signaling with feedback loops

To better understand the signaling activities in the MAPK and AKT regulatory networks, we followed the ISA modeling approach [27] to construct a simplified model of these pathways. We based our starting model (see Figure 1A) on the simplified, joint drug-model of [27]. The starting model includes three receptor tyrosine kinases (RTKs), a simplified representation of the MAPK and AKT pathways, and 10 important oncogenic mutations which are observed in breast cancer cells. In addition, we now added four known feedback signaling events.

Briefly, in ISA, the steady state activity of a signaling molecule is modeled as a latent variable with a continuous value between 0 and 1. The signaling activity is a function of a basal activity, the expression of the signaling molecule itself, the activities of upstream signaling molecules, the effect of mutations, and any kinase inhibitors that are present. Importantly, the parameters of the model are shared across all cell lines and conditions. That is, the parameter values are the same for all cell lines in all conditions. For example, while each cell line can have a different amount of AKT activity in a particular condition, a given amount of AKT signal always gives rise to the same amount of input signal to MTORC1.

To be able to include feedback events in ISA, we make two main changes to the modeling approach. First, to constrain the activity of the signaling molecules between 0 and 1, we use a logistic function rather than a clamping function (see Figure 1B). Although this is computationally more expensive, it makes the activation function smooth, which greatly simplifies solving the systems of equations when feedbacks are present. Second, to calculate the activity of the molecules which are part of a feedback system, we use Newton-Raphson root-finding rather than calculating the activities in one pass from upstream to downstream, as was done before. Since the computation of one Newton-Raphson step scales cubically with the number of signaling molecules, a significant speed-up can be obtained if we restrict the root-finding to the part of the model that contains feedback. We therefore first identify the strongly connected components in the model, using Tarjan’s strong connectivity algorithm [33], illustrated in Figure 1D. This algorithm also provides a topological ordering, which is needed to calculate the signaling activities in the decomposed equations in the correct order. The system of equations corresponding to each strongly connected component is then solved separately, with the benefit that these systems are typically much smaller than the complete model. All equations and the methodology for solving them are described in detail in the Methods section.

### Feedback activity from ERK to RAF is partially identifiable from pre-treatment, non-intervention data

Having a methodology to infer steady state signaling activities in models with feedback, we first wondered whether feedback loops are already identifiable using non-intervention data. To test this, we fitted the starting model (Figure 1A) to the phosphorylation data of thirty, untreated breast cancer cell lines, grown under normal culturing conditions; i.e. the phosphorylation data of Jastrzebski *et al* [27]. We first included only the phosphorylation data of the downstream signaling kinases (Figure 2A). To our surprise, these non-intervention data already suggest that the feedback loop from ERK to RAF is likely to be active (Figure 2B). A possible explanation for the identifiability of feedback from non-intervention data may be that there are many inputs into the MAPK pathway; in the model we included three different RTKs, as well as mutations in KRAS or BRAF. These five inputs together could lead to over-activation of the MAPK pathway, so to prevent this, the inputs need to be restrained in some way. It is easier to accomplish this by a negative feedback loop, since this requires only one parameter to be given a high value, rather than by each input to the MAPK pathway being weak, which requires five parameters to have a low value. In such situations, the Bayesian inference follows the principle of Occam’s razor by preferring a simple explanation over a more complex one.

**Figure 2:**
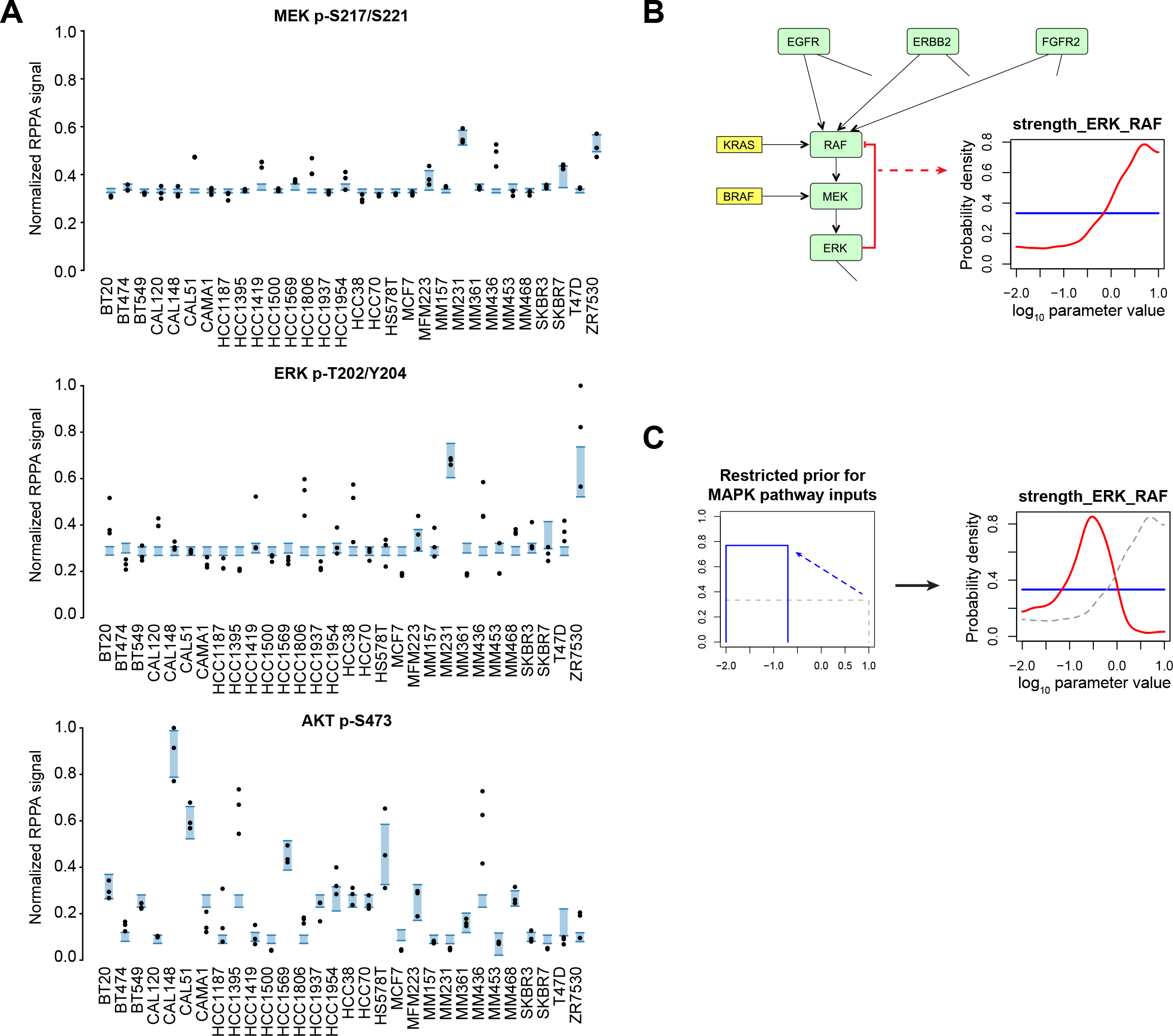
Identification of feedback activity from ERK to RAF from pre-treatment, intracellular signaling measurements. (A) Data and posterior predictive for three of the six epitopes. Black dots indicate the measurement data and the shaded blue area is the 90% confidence interval of the posterior predictive distribution. (B) Posterior probability density for the strength of ERK->RAF feedback signal inferred from the data. (C) Restricting the prior for the inputs to the MAPK pathway to low values results in a weaker RAF->ERK feedback.

To test whether it is indeed the inputs to the MAPK pathway that determine the high values for ERK-to-RAF feedback activity, we artificially forced the inputs to be weak by restricting the prior for the strength parameters of these inputs (Figure 2C). In this case, we indeed see that the feedback becomes weaker as well, indicating that the negative feedback is used to balance the inputs.

Nevertheless, the posterior probability distribution for the strength of the ERK-to-RAF feedback loop has non-zero probability for low values as well. This means that a weak or very weak feedback loop is also consistent with the data. A model entirely excluding this negative feedback loop is less likely to represent the data, but the difference is small (the log Bayes factor is 1.0 in favor of the model including the feedback loop). In summary, given the pre-treatment, non-intervention data of the intracellular kinases, this feedback loop is likely to be active, but based on the single dataset used so far we cannot rule out that it the feedback from ERK to RAF is inactive.

### Adding data can increase the uncertainty in individual parameters

We next tested whether the addition of more data helps in identifying the feedback loop. We first added data for the surface receptor tyrosine kinases, which provide information on the inputs to the pathways (Figure 3A). Interestingly, if we include phosphorylation and protein expression data of EGFR and ERBB2, the feedback activity from ERK to RAF in fact becomes less identifiable (Figure 3B). With these data, the model can infer that the activities of EGFR and ERBB2 are low in the majority of cell lines, and this constrains at least some inputs to the MAPK pathway to low strengths. Given this, a strong feedback is no longer required to balance the activation of the MAPK pathway. Since the feedback is not required, but still feasible, the activity of the feedback has become less certain.

**Figure 3:**
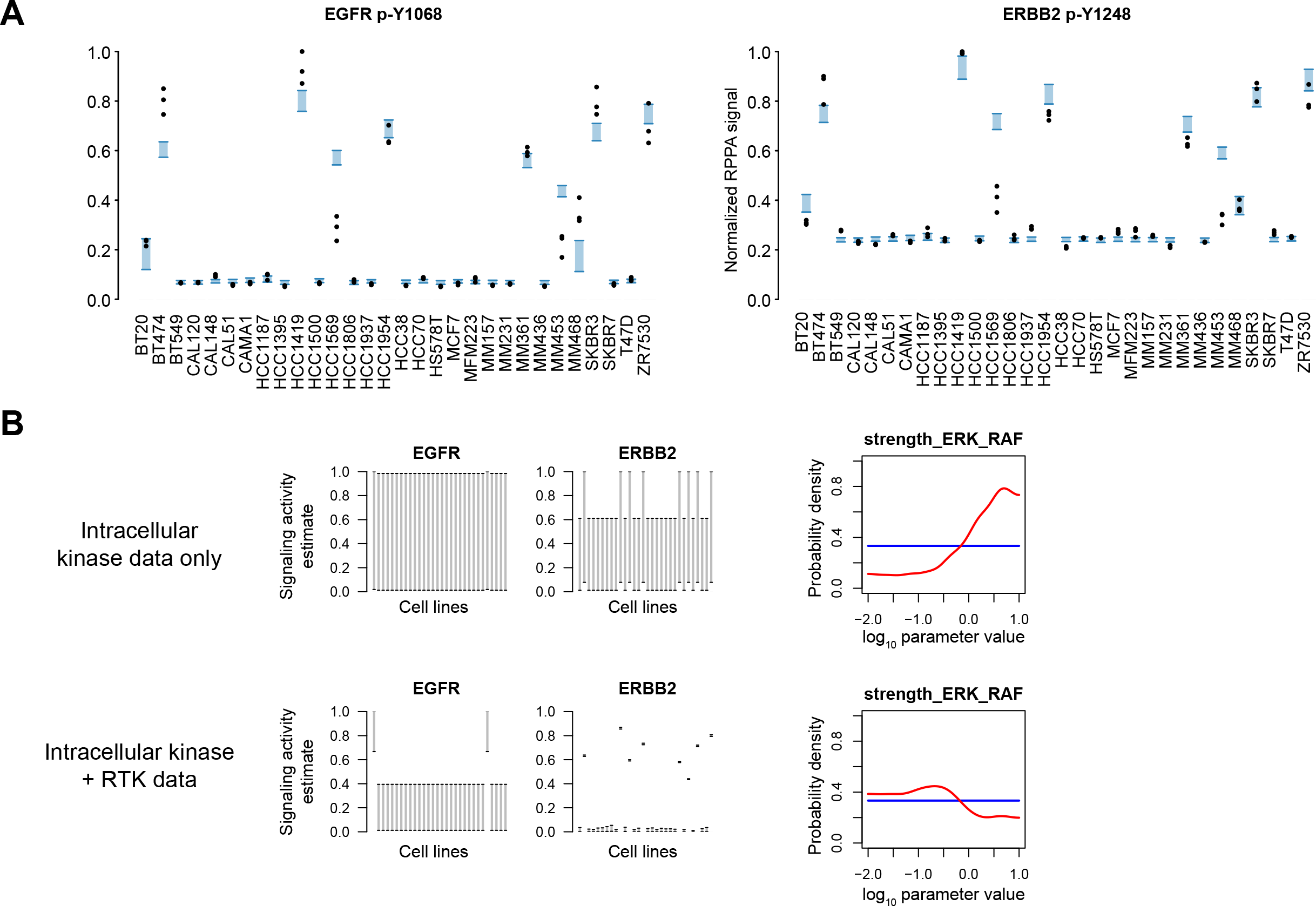
Addition of RTK phosphorylation data increases the uncertainty in ERK to RAF signaling. (A) Data and posterior predictive of the two epitopes that were added. Note that there is cross-reactivity between the antibodies against EGFR p-Y1068 and ERBB2 p-Y1248, hence both measurements are a linear combination of the EGFR and ERBB2 signaling estimates. (B) Posterior probability densities of the ERK->RAF feedback loop, and the signaling estimates of EGFR and ERBB2, with and without the RTK data. For the signaling estimates, the shaded gray areas indicate the 90% confidence interval.

To see whether the RTK data does provide some information overall, we tested whether the joint uncertainty in all parameters does decrease. This can be quantified using Occam’s factors [41], which measure the reduction in the parameter space that is accessible from the posterior compared to the prior. The model with RTK data has a log Occam factor of −127.6, compared to −97.4 without the RTK data. The posterior space collapses more when the RTK data is included than when it is excluded, meaning that the uncertainty in all parameters together is reduced by the inclusion of the RTK data. Together, this shows that, while the total uncertainty is reduced by adding data, the uncertainty of individual parameters can increase.

### Incorporating on-treatment measurements confirms ERK to RAF feedback activity

Intervention data should be more informative for identifying feedback loops. We therefore performed on-treatment measurements, using the MEK-inhibitor PD0325901, the PI3K inhibitor GDC0941 and the dual MTORC1/2 inhibitor AZD8055, in a smaller panel of cell lines (Figure 4A). Cells were either allowed to grow exponentially for 48 hours, or were starved for 24 hours, followed by 30 minute pre-treatment with inhibitors, before stimulation with EGF to increase signaling activity. The EGF stimulation is included in the model by setting EGF activity to 1 for the EGF-stimulated conditions and to 0 otherwise.

**Figure 4:**
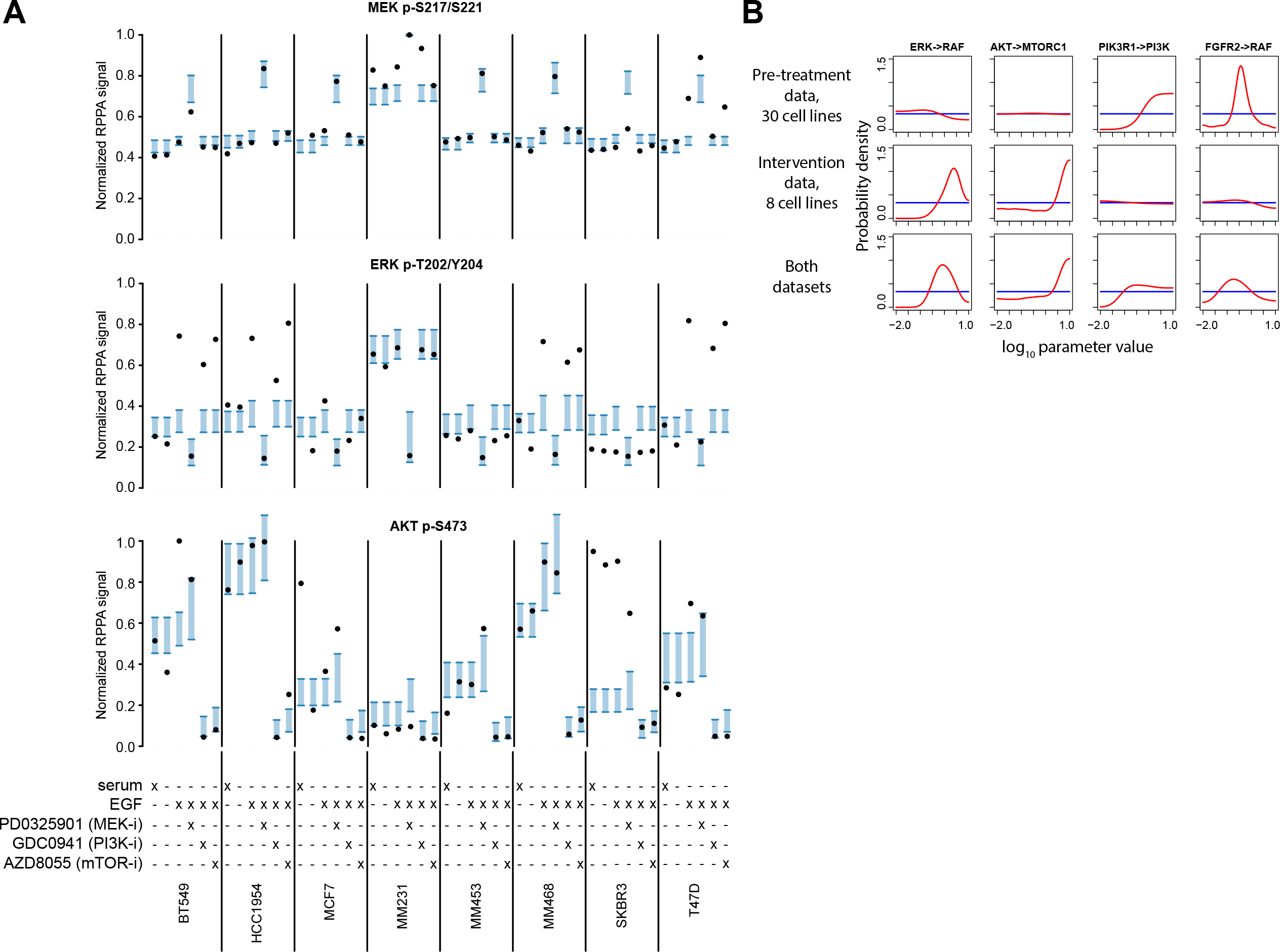
On-treatment data constrains the feedback from ERK to RAF. (A) Data and posterior predictive for three of the eight epitope added to the inference. The posterior predictive is obtained from inference with the pre-treatment and intervention datasets combined. (B) Comparison of densities of the ERK->RAF feedback loop and three other parameters with either dataset alone or together.

This on-treatment dataset was generated separately, at a different time, from the 30-cell line pre-treatment dataset. Although the RPPA measurements include a set of standard control lysates, the spot intensities are quantified separately and there are likely to be differences between the two data batches. Furthermore, we cannot always rely on a sufficient number of overlapping measurement conditions to align multiple datasets. We therefore incorporated batch correction directly into the inference, such that the differences between batches are automatically accounted for, and balanced against the most likely signaling strengths. This is accomplished by adding an offset and a scale parameter for each epitope in each dataset (see Figure 1C and the Methods section for details).

Focusing on the MEK inhibitor, we see that using the on-treatment data, feedback activity from ERK to RAF is clearly identifiable (Figure 4B, first column). This shows that on-treatment data is indeed more informative for inferring feedback activity, as expected. The result of the feedback is also clearly visible in the data (Figure 4A) given that MEK phosphorylation greatly increases after treatment with a MEK inhibitor. Although the increase in MEK phosphorylation upon MEK inhibitor treatment could also be a direct effect of the inhibitor rather than a feedback event (discussed in more detail for ATP-competitive AKT inhibitors later), PD0325901 is an allosteric inhibitor [43,44], and the increased phosphorylation of MEK is rather more likely to be through ERK to RAF feedback signaling [45].

When considering several other parameters, we see that both the pre-treatment and on-treatment measurement provide useful information (Figure 4B). The pre-treatment measurements were collected for a single condition for more cell lines, while the on-treatment measurements included more conditions for fewer cell lines. Combining the two datasets provides a broad coverage over cell lines as well as intervention measurements in a smaller panel to help identify signal strengths. The on-treatment data not only helps to identify feedback activity from ERK to RAF, but also, for example, the strength of AKT to MTORC1 signaling. Conversely, the broader pre-treatment data helps identify, for example, the effect of PIK3R1 mutations and FGFR2 amplifications, neither of which were found in the smaller cell line panel used for generating the on-treatment data.

### Negative feedback is likely to be active in the AKT pathway

One of the main advantages of f-ISA lies in identifying which signaling activities and strengths are most strongly supported by the data. This advantage becomes most apparent when there are multiple interrelated feedback events, combined with information from multiple datasets. In this case it is impossible to manually keep track of all the constraints, and computational modeling is necessary to obtain a quantitative understanding of the regulatory network. To illustrate this, we next focus on the AKT pathway, where several different feedback events exist, further complicated by conflicting reports in the literature. As a first approximation, and as can be seen in Figure 1A, we incorporated the two hypotheses regarding feedback through SIN1, described in the introduction, through a negative link between MTORC1 and MTORC2 (representing inhibitory phosphorylation of SIN1 by S6K), and a positive link between AKT and MTORC2 (representing stimulating phosphorylation of SIN1 by AKT). The feedback through IRS1 is included as a negative link between MTORC1 and PI3K.

Using the two datasets described so far (see Figure 4A and 5A), the model identifies that there is likely to be a negative feedback in the AKT pathway (Figure 5B). The positive feedback through SIN1 is identified to be weak (Figure 5B, top-left panel), while the negative feedback through SIN1 is likely to be active (Figure 5B, top-right panel). This is partially driven by the same effect as in the MAPK pathway, that is, there are again many inputs into the AKT pathway, which are most easily balanced by a negative feedback loop. This effect can also be seen by a very strong correlation between the strength of signaling from PI3K to MTORC2 and the negative feedback from MTORC1 to MTORC2 (Figure 5B, bottom-right panel). However, apart from this effect, the model also infers that the negative feedback from MTORC1 to MTORC2 is used to reduce AKT-pathway activity in cell lines with high MAPK pathway activity, including MDA-MB-231 and ZR-75-30.

**Figure 5:**
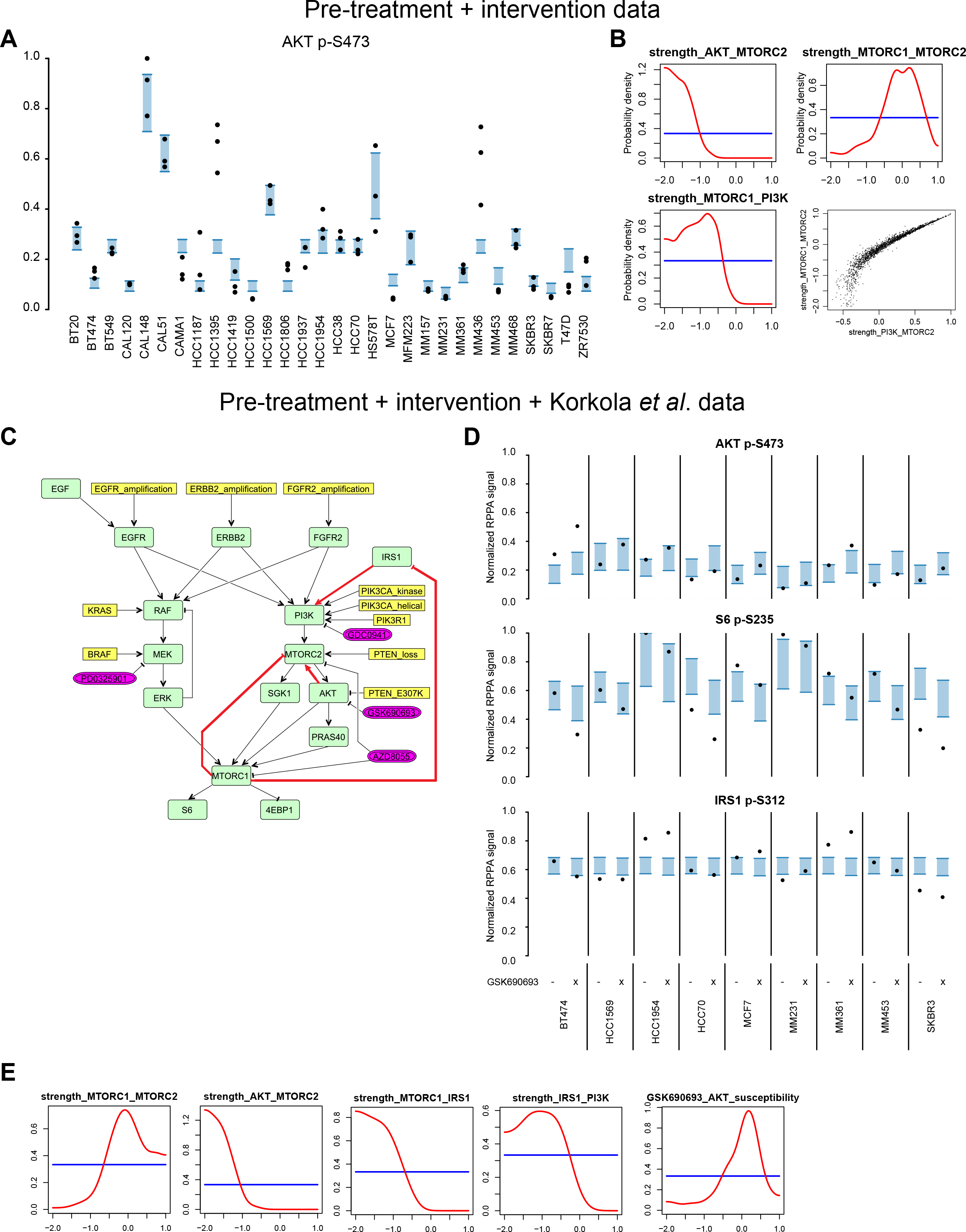
Feedback in the AKT pathway is likely to be negative. (A) Data and posterior predictive of AKT p-S473 in the pre-treatment dataset in the inference with the pre-treatment and intervention datasets combined; c.f. Figure 2A where the same data is shown but the intervention dataset was not included. (B) Feedback signaling strengths in the AKT pathway. The bottom right panel shows a scatter plot of the samples of the PI3K->MTORC2 and MTORC1->MTORC2 signal strengths. (C) Model with GSK-690693 as an additional inhibitor, and IRS1 explicitly included as a signaling molecule. The links for which the posterior is shown in E are highlighted in red. (D) Data and posterior predictive for three of the epitopes used for the inference; the data is the 1-hr time point of the dataset of Korkola *et al* [42]. The previous two datasets are included in the inference as well. (E) Posterior distribution of the feedback signaling events in the AKT pathway, given the three datasets together.

The negative feedback in the AKT pathway could be achieved in several different ways, either through IRS1 or through SIN1. To further resolve which of these proteins mediates the feedback, we extended the model to explicitly include IRS1 as signaling molecule (Figure 5C), and we incorporated information from the dataset of Korkola *et al* [42], who performed time-course RPPA measurements with a different panel of cell lines upon inhibitor treatment, including the AKT inhibitor GSK690693. We selected the 1 hour time point from their dataset as it is closest to the 50 minute inhibitor treatment (30 minute inhibitor pre-treatment plus 20 minute subsequent EGF stimulation) in our intervention dataset. To accommodate this dataset, we also added GSK690693 as inhibitor to the model.

A complication when using ATP-competitive AKT inhibitors like GSK690693 is that these inhibitors can directly alter the susceptibility of AKT phosphorylation by PDK1 and MTORC2, independently of AKT kinase activity [21]. This effect causes an increased AKT phosphorylation upon AKT inhibitor treatment, independent of any feedback events. We incorporated this effect in the model by allowing a kinase inhibitor to alter the strength of incoming signals to the targeted protein (see Methods section; note that this effect cannot be seen in the SBGN schematics but is included in the SBML model annotation).

The model can fit several aspects of the Korkola data, including the expected reduction in S6 phosphorylation and increased AKT phosphorylation upon AKT inhibitor treatment (Figure 5D), without compromising the fit of the other two datasets (Supplementary File 1). Furthermore, we see that the increased AKT phosphorylation upon AKT inhibitor treatment is indeed most likely explained by an increase in AKT phosphorylation susceptibility (Figure 5E, right-most panel). Despite this, a negative feedback in the AKT pathway is still likely to be present (Figure 5E, left-most panel). This, however, is unlikely to be through IRS1 S312 phosphorylation (Figure 5E, 4^th^ and 5^th^ panel). Consistent with this, the measured IRS1 S312 phosphorylation levels are not consistently changed upon AKT inhibitor treatment in this dataset (Figure 5D, bottom panel). Finally, the presence of a positive feedback in the AKT pathway, acting via SIN1, is still deemed unlikely in this model with IRS1 included as signaling molecule (Figure 5E, 2^nd^ panel).

### Testing the strength of negative feedback through IRS1 by modulating its expression

The above described inference indicated that while there is negative feedback in the AKT pathway, the feedback through IRS1 S312 phosphorylation is likely to be weak. However, there is uncertainty in the strength of IRS1 to PI3K signaling. To further resolve this, we experimentally disrupted the IRS1 feedback mechanism by pre-treating cells with the IGF1R inhibitor NT157, which has been shown to induce degradation of IRS1 [46,47], and measured how this affects phosphorylation upon AKT- or S6K inhibitor treatment (Figure 6A). To accommodate this data more precisely, it is also useful to expand the model by including S6K and SIN1 as separate nodes (the resulting model is shown Figure 6B). Since we now explicitly include SIN1, we model the unknown effect of SIN1 on MTORC2 by allowing this signal to be either positive or negative.

**Figure 6:**
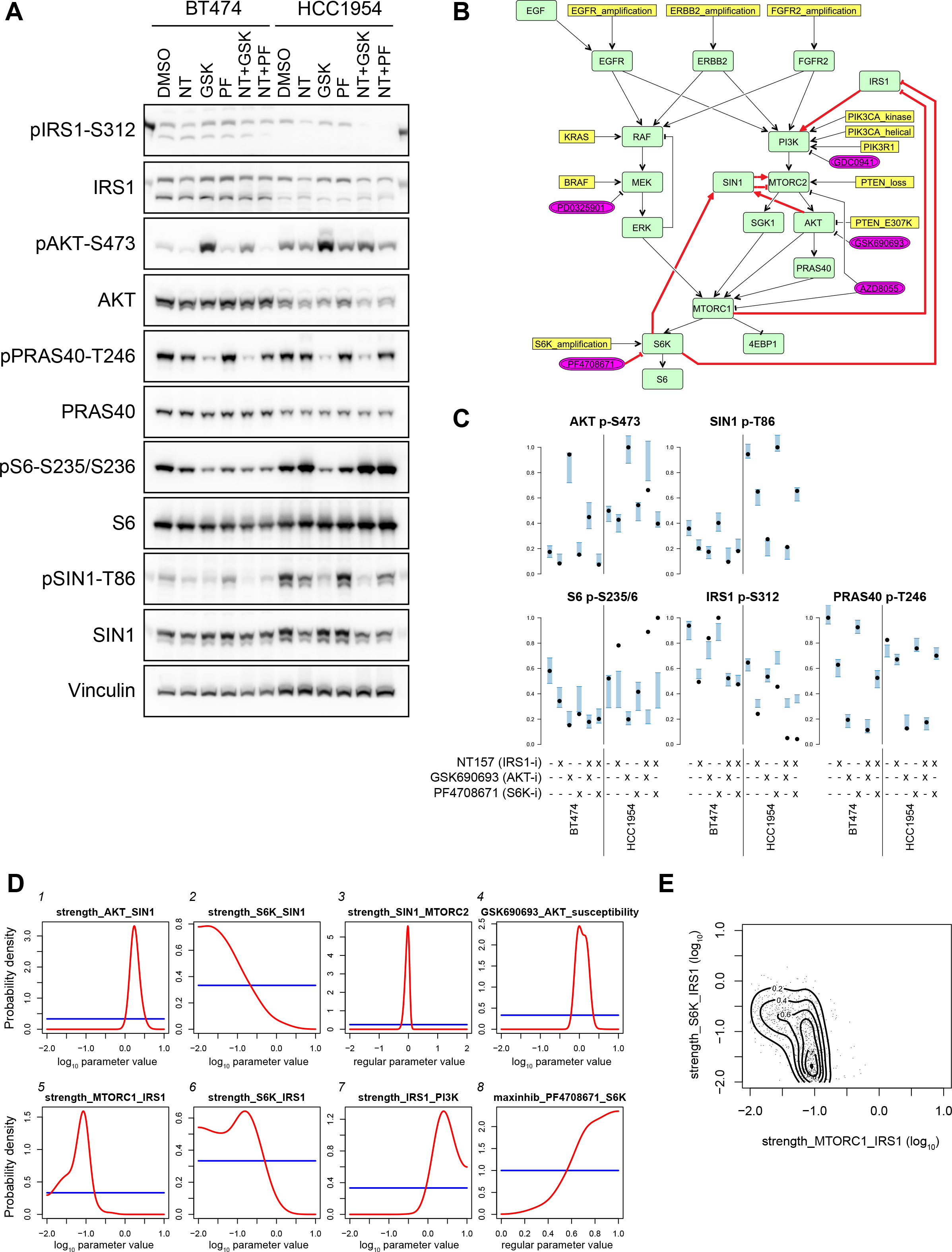
Resolving the feedback through IRS1. (A) BT474 and HCC1954 cell lines treated with the IRS1/2 inhibitor, NT-157 (NT), alone or in combination with either the AKT inhibitor, GSK690693 (GSK), or the S6K inhibitor, PF4708671 (PF), were analyzed by immunoblotting for expression and phosphorylation of key molecules in the AKT pathway. Control samples were treated with DMSO alone. (B) Model with S6K, SIN1 and PF4708671 added. The double, dashed arrows from SIN1 and MTORC2 indicate that the sign of this link is unknown. The links for which the posterior is shown in D are highlighted in red. (C) Quantification of the western blot signals shown in A, along with the posterior predictive of the fitted model. The previous three datasets are included in the inference as well. (D) Posterior distribution of the feedback signaling strengths in the AKT pathway, given the four datasets together. (E) Scatter plot of the posterior samples for the MTORC1->IRS1 and S6K->IRS1 signal strengths, with a 2D-KDE of the bivariate marginal posterior shown in contours.

With some exceptions, the model can describe the IRS1-disruption experiment very well (Figure 6C), without compromising the fit to the other datasets (Supplementary File 1). The data shows a clear increase in AKT phosphorylation after AKT-inhibitor treatment, but not after S6K-inhibitor treatment. We also see that the increased AKT phosphorylation is reduced by pre-treatment with NT157. This is consistent with the report that AKT-inhibitor-induced AKT-phosphorylation is dependent on MTORC2 activity [21], and MTORC2 being dependent on IRS1 signaling. The S6K inhibitor, although it does reduce S6 S235/236 phosphorylation, does not change the phosphorylation levels of AKT S473, SIN1 T86 or IRS1 S312, suggesting it is not critically involved in mediating the feedback loop under these conditions.

We can then use the model to delineate all the feedback activities (Figure 6D). In contrast to the inference based on the previous datasets, the model can now infer that the signal from IRS1 to PI3K is strong, perhaps even a point of signal amplification in the pathway, given that the inferred strength is larger than 1 (Figure 6D, panel 7). Hence a relatively weak negative feedback to IRS1 can still affect the AKT pathway. Furthermore, there appears to be a weak, though nonzero, feedback to IRS1 (Figure 6E). The model cannot entirely resolve whether the feedback is mediated by MTORC1 only or also through S6K (Figure 6E), due to the uncertainty in how strongly the S6K inhibitor is inhibiting its target (Figure 6D, panel 8) - the more likely it is that PF4708671 is inhibiting S6K, the less likely S6K is to phosphorylate IRS1 on S312. Despite this uncertainty, an S6K-independent signal to IRS1-pS312 is approximately three times more likely than an S6K-dependent signal (Figure 6D and E).

The hyperphosphorylation of AKT upon GSK690693 treatment is still mainly explained by the inhibitor-induced change in phosphorylation susceptibility (Figure 6D, panel 4). It is also clear that SIN1 T86 phosphorylation is dependent on AKT (panel 1), providing further information for the debate on the regulation of this site. Furthermore, SIN1 T86 phosphorylation is unlikely to be mediated by S6K (panel 2), although there is again some amount of uncertainty caused by the poor and uncertain efficacy of the S6K inhibitor (panel 8). However, given the present data, feedback through SIN1 is predicted to have only a minor negative effect on MTORC2 activity, if any (panel 3).

Taking into account 1254 data points spread across 108 different conditions, the most likely feedback in the AKT pathway is a strong effect of AKT back to SIN1, which only weakly inhibits MTORC2, combined with a weak signal from MTORC1 to IRS1 S312 (but unlikely through S6K), which is however amplified by IRS1.

### Using ISA to test the agreement between datasets

Individual datasets are often insufficient to constrain most of the signaling strengths. Figure 7A shows how much each parameter is constrained by each dataset, as well as all datasets together. We can see that in many cases, a parameter is constrained by a single dataset. In this way, each dataset provides complementary information, and thus all datasets together are able to constrain most of the parameters to some extent.

**Figure 7:**
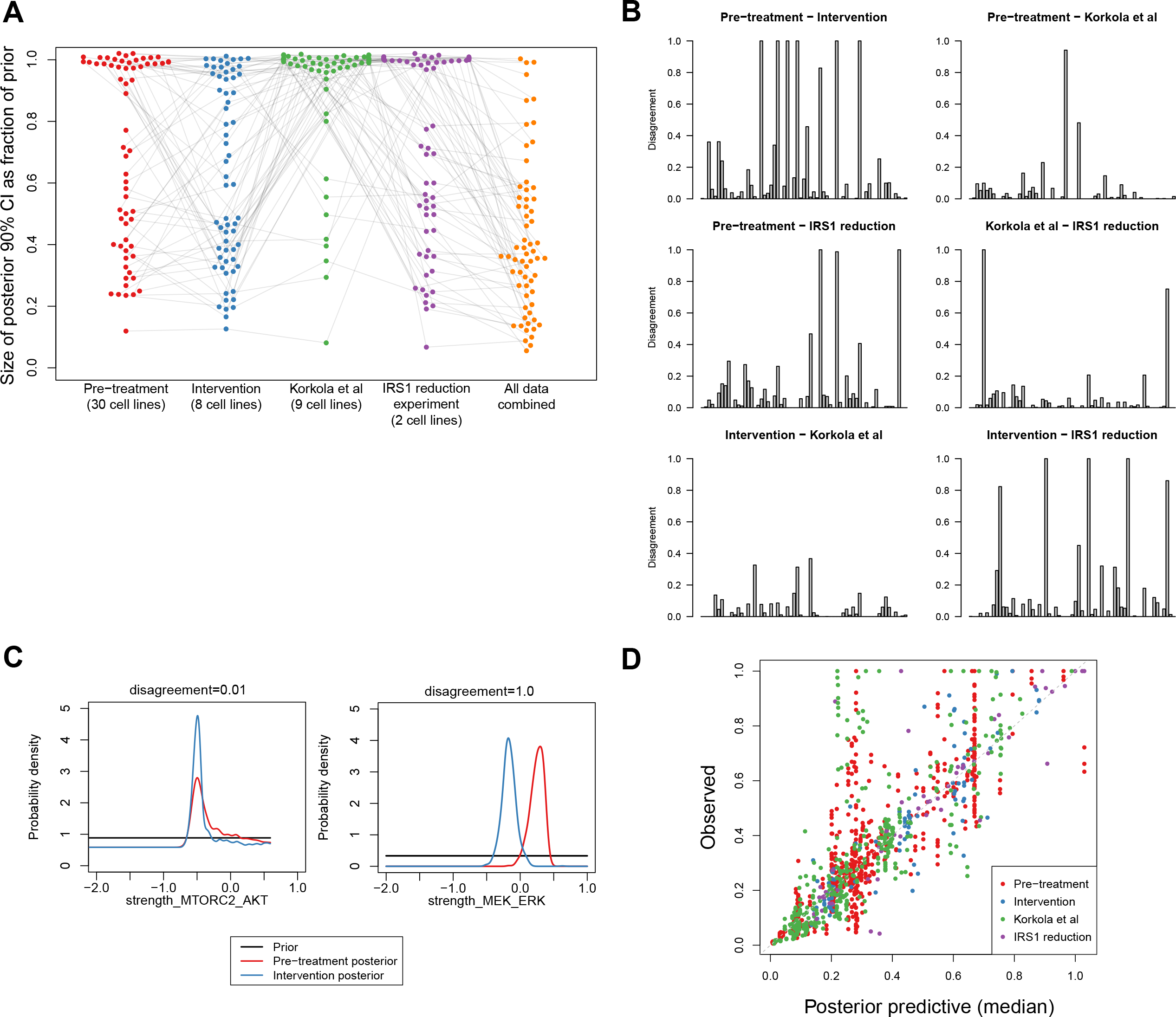
Comparing the model fit and estimates across datasets. (A) Reduction in uncertainty for each of the 63 signaling parameters, given each dataset separately and together. Each dot indicates one parameter. Grey lines connect the same parameters between datasets. Some parameters are constrained by multiple datasets, while other parameters are uniquely constrained by one dataset. (B) Disagreement in parameter estimates between datasets; each bar represents on of the 63 signaling parameters. Disagreement is calculated as one minus the overlap coefficient, that is, 1 − *IS* / min(*CI*_1_, *CI*_2_), where *CI*_*i*_ is size of the posterior 90% confidence interval given each dataset, and *IS* is the size of the intersection of the two CIs. Detailed figures with labels are included as Supplementary Figure 1. (C) Examples of two parameters for which the pretreatment dataset and the intervention dataset either agree (top panel) or disagree (bottom panel). (D) Scatter plot of all fitted data, represented by the median of the posterior predictive distribution, against the observed values.

Since datasets often provide different measurements, it is not clear how they can be directly compared against each other. However, we can use f-ISA to test whether datasets agree with each other given a model. To explore this, we calculated the disagreement in each parameter between any pair of the four datasets. We calculate the disagreement by taking one minus the overlap coefficient between the posterior 90% confidence intervals (CI) of the parameter given each dataset, where the overlap coefficients is the size of the intersection divided by minimum size of either CI. A disagreement of 1 means that the posterior 90% CIs do not overlap, whereas a disagreement of 0 means that the CI of one posterior is completely contained within the other, or are even exactly the same. Figure 7B shows this disagreement between datasets. Reassuringly, for most of the parameters the disagreement is small, indicating that in most cases the posterior distribution is accurate between datasets (an example is shown in Figure 7C). Note that this also includes many cases where one or both of the datasets simply do not constrain a parameter at all, hence the model may not be able to give a precise prediction for a parameter based on a dataset, but the posterior is still accurate in that it quantifies the uncertainty correctly.

There are, however, also some parameters for which the datasets disagree. Most disagreement occurs between the pre-treatment and intervention datasets. For example, the strength of signaling from MEK to ERK is estimated differently by the pre-treatment and intervention datasets. Assuming that the measurements in both datasets can be trusted, this would indicate that the model is overestimating its certainty. Nevertheless, despite the disagreement in some parameters, the model can provide a good fit against all data simultaneously (Figure 7D), and the resulting joint posterior is the most likely compromise between the posteriors inferred by the datasets separately.

### Identification of unexplained data points

Such models can also be used to identify unexpected parts of the data. Although the model can explain a large part of the data (Figure 7D), there are also data points which cannot be recapitulated by the model. For example, in the intervention dataset, EGF stimulation leads to a large increase in ERK phosphorylation in four of the eight cell lines, which the model cannot reproduce (see Figure 4A). The cell line specificity is partially explained by EGFR expression, but not completely, given that despite EGFR expression is included as input the model cannot explain the variability observed. In addition to the cell line specificity, there is only a very modest increase in MEK phosphorylation upon EGF stimulation, raising the question of how the signal from EGF to ERK is transmitted. One explanation for a difference in the changes of MEK and ERK phosphorylation could be that MEK amplifies the signal. However, the activation function allows for a signal amplification of up to 10-fold, and hence the constraints posed by all other data do not seem to allow this explanation. An explanation rooted in signal amplification also does not explain why this occurs only in some of the cell lines. Alternative explanations might be that there are additional feedback mechanisms or feed-forward signaling pathways which transmit the EGF signal downstream, that other phosphorylation sites are involved, or that the discrepancy is a result of the specific timing of starvation, inhibition and stimulation.

Another interesting part of the data that cannot be explained by the model, is the increased S6 phosphorylation in HCC1954 upon NT-157 pretreatment (Figure 6C). It has been shown that NT-157 treatment can induce MAPK pathway activation by inducing IGF1R-SHC complex formation [46], and the increased MAPK signaling could lead to elevated S6 phosphorylation [48,49]. These are links that could be included in the model, although it would then still be unclear why the increased S6 phosphorylation is specific for HCC1954 and does not occur in the BT474 cell line. Alternatively, other pathways may be activated as a result of the disruption of IRS1 signaling, such as a stress response, potentially leading to S6K activation. Indeed, following treatment with NT-157, we did observe a reduction in cell viability of the HCC1954 cell line, while the BT474 cells appeared unaffected by the treatment.

These two examples show that the computational model can highlight cases where our knowledge, as summarized in the model, is incomplete and unable to fully describe the data. To resolve these discrepancies, additional rounds of model extension, combined with further experimental measurements to constrain the parameters, can be performed. Alternatively, the identified discrepancies can help define precise questions for follow-up functional genetic screening.

## Discussion

As our knowledge of biological signaling networks grows, it is increasingly difficult to fully understand the behavior of these networks. Computational modeling then provides a means to quantify the interactions as well as the contributions of different signaling molecules and genetic aberrations. We can also use these models to test whether our knowledge is sufficient to explain the data. As our models of biological signaling get more complex, we will also increasingly need to integrate multiple datasets to be able to identify the key parameters.

In this work, we used four datasets to constrain the parameters in a signaling model. Each of the data sets provided complementary information, and the largest number of the parameters could only be constrained with all four datasets employed simultaneously. We also described several non-intuitive behaviors of the identifiability of parameters. We found that feedback loops can sometimes be partially identified from non-intervention data, including the negative feedback from ERK to RAF, as well as a feedback from MTORC2 to MTORC1 used to reduce AKT signaling activity in cell lines with high MAPK pathway activity. Additionally, for the feedback from ERK to RAF, we found that adding more data to the inference can lead to an increased uncertainty in individual parameters. This is not caused by inconsistency of the data but by shifts in which signaling activities are more likely to explain all data simultaneously. We find that the aggregate uncertainty in all parameters does decrease when more data is employed in the inference.

Various other modeling approaches have previously been used to understand these signaling pathways. Thobe *et al* used logical modeling of MTORC2 signaling, including hypotheses of the regulation through SIN1 [50]. However, they were not able to resolve a preference between models, and given such a non-quantitative approach all models were found to be in agreement with the data. We find that using a quantitative approach as presented here, it is possible to deduce a preference for one feedback loop over another. Dalle Pezze *et al* constructed ODE models of mTOR signaling [51], although they did not include the SIN1 feedbacks (these had not been reported yet). In their ODE model, they used fixed parameter values, although they did perform a sensitivity analysis around the optimum. Given that in our study there are 108 model simulations required to evaluate the model with respect to the four datasets, and the many parameter values that need to be considered to characterize the multidimensional parameter space, it is presently not feasible to use such detailed ODE models in combination with a full uncertainty analysis. Nevertheless, with a quasi-steady state approach we were able to quantify the uncertainty in the feedback mechanisms.

Alongside the abovementioned advantages, the f-ISA modeling framework posseses several disadvantages as well. First, the model is restricted to known mechanisms. It is possible that other links provide an equally good fit. For example, there could be other negative feedbacks ending in or upstream of MEK and AKT, or the involvement of other phosphorylation sites on IRS1. Our approach does not search for additional links that may fit the data better. Rather, the goal is to test whether a particular model can describe the data well. Unfortunately, combining extensive parameter uncertainty analysis with a search for network topology is computationally intractable at present. Second, the calculation of signaling with feedback events as done here is significantly slower than the original ISA modeling approach without feedback. It is therefore still impractical to infer signaling activities for large models with feedback while also including full dose response curves, especially with multiple drugs. For the dose response data presented in Jastrzebski *et al* [27], this would add another 300 model conditions per drug, as well as additional model parameters. It may be feasible to perform inference sequentially, thereby reducing the number of model evaluations required [52]. Alternatively, evaluating the gradient of the likelihood and using sampling algorithms that can leverage this gradient information may speed up the inference [53,54].

A third limitation is that f-ISA is currently only able to handle feedback events which occur on one timescale. We focused here on fast-acting post-translational feedback. This approach could also be used to study transcriptional regulation; in this case total protein or mRNA levels could be used as inference data instead of phosphorylation data. It is also possible to include both transcriptional and post-translational feedback in the same model, but this would assume that the feedbacks occur on approximately the same timescale, which is probably unreasonable. To disentangle feedback mechanisms acting over different timeframes, it would be necessary to add further extensions that can calculate steady states with feedback loops occurring on multiple timescales, or it may be more beneficial to switch to dynamic models in this case.

Despite these limitations, we believe f-ISA is a useful approach to delineate context-dependent activities in signaling networks with multiple feedback paths. By iteratively incorporating additional detail in the signaling networks, and additional measurements to constrain the parameters, it is possible to obtain an increasingly thorough, quantitative understanding of a regulatory signaling network.

## Acknowledgments

We are grateful to Jordi Vidal Rodriguez for experimental support with the on-treatment, intervention RPPA experiment.

